# Engineered probiotics for local tumor delivery of checkpoint blockade nanobodies

**DOI:** 10.1101/562785

**Authors:** Candice Gurbatri, Courtney Coker, Taylor E. Hinchliffe, Ioana Lia, Samuel Castro, Piper M. Treuting, Nicholas Arpaia, Tal Danino

## Abstract

Immunotherapies such as checkpoint inhibitors have revolutionized cancer therapy yet lead to a multitude of immune-related adverse events, suggesting the need for more targeted delivery systems. Due to their preferential colonization of tumors and advances in engineering capabilities from synthetic biology, microbes are a natural platform for the local delivery of cancer therapeutics. Here, we present an engineered probiotic bacteria system for the controlled production and release of novel immune checkpoint targeting nanobodies from within tumors. Specifically, we engineered genetic lysis circuit variants to effectively release nanobodies and safely control bacteria populations. To maximize therapeutic efficacy of the system, we used computational modeling coupled with experimental validation of circuit dynamics and found that lower copy number variants provide optimal nanobody release. Thus, we subsequently integrated the lysis circuit operon into the genome of a probiotic *E. coli* Nissle 1917, and confirmed lysis dynamics in a syngeneic mouse model using *in vivo* bioluminescent imaging. Expressing a nanobody against PD-L1 in this strain demonstrated enhanced efficacy compared to a plasmid-based lysing variant, and similar efficacy to a clinically relevant monoclonal antibody against PD-L1. Expanding upon this therapeutic platform, we produced a nanobody against cytotoxic T-lymphocyte associated protein -4 (CTLA-4), which reduced growth rate or completely cleared tumors when combined with a probiotically-expressed PD-L1 nanobody in multiple syngeneic mouse models. Together, these results demonstrate that our engineered probiotic system combines innovations in synthetic biology and immunotherapy to improve upon the delivery of checkpoint inhibitors.

**SENTENCE SUMMARY:** We designed a probiotic platform to locally deliver checkpoint blockade nanobodies to tumors using a controlled lysing mechanism for therapeutic release.

## INTRODUCTION

Cancers exploit checkpoint signaling pathways by expressing ligands such as programmed cell death protein-ligand 1, PD-L1, which bind to the PD-1 receptor to inhibit the activation, expansion, and function of T cells and support immune evasion (1–3). These inhibitory checkpoint blockade mechanisms have prompted the exploration of blocking monoclonal antibodies (mAbs) as therapeutics against these molecules. While anti-PD-L1 mAbs have achieved some level of tumor regression in ~30% of cancers like melanoma (4, 5), they can also result in immune-related adverse effects (iRAEs), with up to 70% of patients experiencing a range of toxicity grades resulting in fatigue, skin rashes, endocrine disorders, or hepatic toxicities (6–9). Furthermore, combination therapies of anti-PD-L1/PD-1 mAbs and anti-cytotoxic T-lymphocyte associated protein-4 (CTLA-4) mAbs are more efficacious than monotherapies, but cause higher grade toxicities that lead to favoring of less efficacious monotherapies or eventual drug discontinuation (10, 11). Moreover, iRAEs are often treated with steroids or other immunosuppressants that may comprise the antitumor response and subsequently affect the efficacy of checkpoint blockade antibodies (9, 12). Thus, there is a clear need for improved delivery of checkpoint blockade inhibitors to the tumor site to minimize adverse effects and shift the treatment preference towards combination therapy approaches.

Rapid development of genetic technologies has enabled the engineering of intelligent microbial delivery systems for therapeutic applications. Specifically, synthetic biology has generated numerous examples of genetic circuits controlling bacteria growth and gene expression (13–19), allowing them to sense and respond to disease states of inflammation, infection, and cancer (20–23). Particularly for cancer, a multitude of studies have shown that systemic administration of bacteria results in their selective colonization of tumors, providing a unique opportunity for tumor drug delivery. This occurs primarily due to reduced immune surveillance along with the ability of bacteria to grow within the hypoxic and necrotic tumor core (24–27). At the same time, microbiome research efforts have revealed the widespread prevalence of microbes within malignant tissue that do not cause infections or other long-term detrimental health effects (28, 29). Since bacteria are both inherently present and selectively grow in tumors, they provide a natural platform for the development of programmable therapeutic delivery vehicles.

Harnessing the converging advancements in both immunotherapy and synthetic biology, we engineered probiotic bacteria to locally and controllably release PD-L1 and CTLA-4 antagonists in the form of blocking nanobodies. Specifically, we coupled immunotherapeutic expression with an optimized lysing circuit mechanism, such that probiotic bacteria carrying the anti-PD-L1 nanobody homes to the necrotic tumor core, grow to a critical density, and lyse effectively releasing the anti-PD-L1 nanobody to block the PD-1/PD-L1 interaction between tumor and T cells (Fig. 1a).

**Figure 1:**
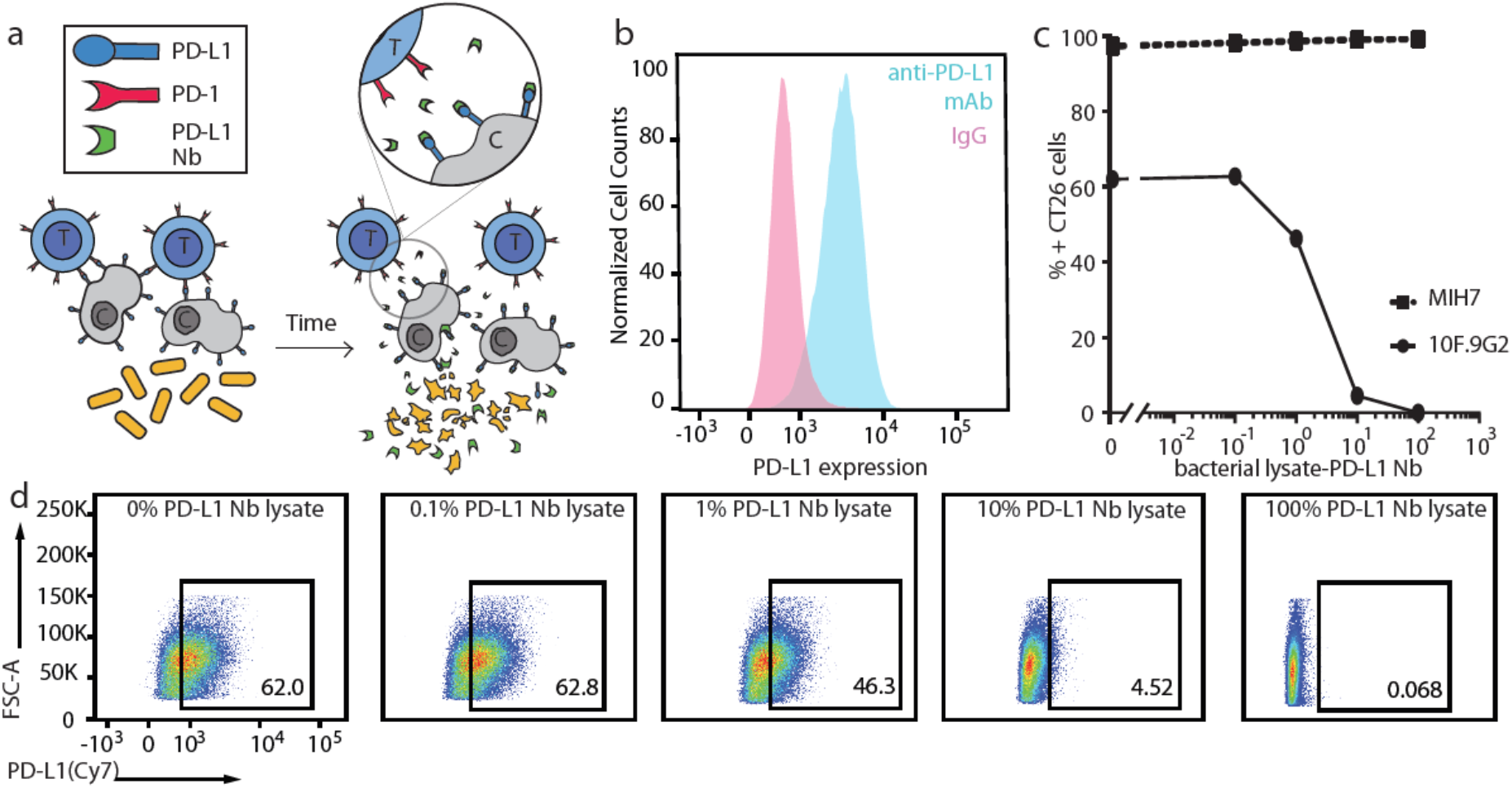
Design of a probiotic cancer therapy system for release of a functional anti-PD-L1 blocking nanobody. **(a)** Bacteria engineered to controllably release a constitutively produced PD-L1 blocking nanobody (Nb), which binds to PD-L1 on the tumor cell surface and blocks its interaction with PD-1 expressed on T cells. **(b)** Flow cytometry analysis of PD-L1 expression on CT26 cells (pink peak: isotype control; blue peak: stained with an anti-PD-L1 mAb, 10F.9G2 clone). **(c)** Binding curve of PD-L1 Nb to the 10F.9G2 and MIH7 PD-L1 epitopes on CT26 cells. (d) Flow cytometry plots of CT26 populations, where CT26 cells were co-incubated with a constant concentration of PE/Cy7-conjugated 10F.9G2 mAb and varying concentrations of bacterial lysates containing the PD-L1 Nb.

## RESULTS

### Construction and characterization of a probiotically-expressed PD-L1 Nb

A single-domain antibody, or nanobody (Nb), blocking PD-L1 was chosen from the RCSB Protein Data Bank as therapeutic cargo. Unlike antibodies with a molecular size of approximately 150 kDa, Nbs are only 15 kDa and lack an Fc region, which requires glycosylation by mammalian cells, and can therefore be produced recombinantly in bacteria (30, 31). Nbs provide multiple advantages, including their small size, which allows for increased diffusion within the tumor microenvironment and more rapid clearance from the bloodstream through glomerular filtration thereby reducing systemic toxicity (32). While faster blood clearance may suggest shorter therapeutic impact, the bacterial platform allows for continuous and intratumoral Nb production to improve upon this limitation. Therefore, the PD-L1 Nb sequence was cloned downstream of a strong constitutive pTac promoter on a stabilized, high copy ColE1 origin of replication plasmid to allow for maximal gene expression. A human influenza hemagglutinin (HA) protein tag was added to 3’ end of the Nb for *in vitro* visualization and an Axe/Txe stability mechanism was cloned into the vector to prevent plasmid loss during bacterial replication (Fig. S1a) (33). The plasmid was transformed into the probiotic strain, *E. coli* Nissle 1917, containing a genomically integrated *luxCDABE* cassette for bacterial tracking *in vivo* (EcN-lux). *E. coli* Nissle 1917 was chosen as the therapeutic vehicle because of its proven safety, as it is currently prescribed for oral administration in humans, and for its ease in genetic manipulation (34).

Flow cytometry analysis (Fig. 1b) and immunofluorescence (Fig. S1b) were used to confirm PD-L1 expression on CT26 cells, a colorectal cell line that has a modest antitumor response to PD-L1 mAb (35, 36). To investigate the binding pattern of a previously uncharacterized PD-L1 Nb, bacterial lysate containing the probiotically-produced PD-L1 Nb was incubated on a monolayer of CT26 cells and an anti-HA mAb was used to probe for PD-L1 Nb binding. Imaging using fluorescence microscopy revealed a similar expression pattern to that of PD-L1 expression probed for by an anti-PD-L1 mAb (Fig. S1b). To quantify binding kinetics of the PD-L1 Nb, multiple dilutions of bacterial lysate containing the PD-L1 Nb were co-incubated on a monolayer of CT26 cells with a constant concentration of fluorescently-tagged anti-PD-L1 mAbs known to bind to either the 10F.9G2 or MIH7 epitope. With a more diluted lysate concentration, an increase in the 10F.9G2 mAb fluorescence was observed (Fig. 1c, d). Notably, a very small fraction – approximately 5%--of the total lysate produced was needed to achieve binding to 50% of PD-L1+ CT26 cells. However, no change in fluorescence of the MIH7 mAb was observed as a function of bacterial lysate (Fig. 1c), suggesting that the PD-L1 Nb specifically binds to an epitope similar to that recognized by 10F.9G2.

### Optimization of lysis circuits for controllable therapeutic release

Since we observed dose-dependent binding of the Nb to PD-L1 on CT26 cells, we sought to enhance therapeutic release by optimizing a synchronized lysis circuit (SLC), whereby a bacterial population lyses once a critical density is reached effectively releasing its therapeutic payload (21). The SLC has been shown to aid in tumor-selective bacterial production, population limitation, and therapeutic release, serving multiple purposes critical for translational efforts. However, one drawback of the original SLC system is its reliance on plasmids. Since the quorum sensing genes and more importantly the lysis gene are cloned onto plasmid vectors, they can be lost or mutated during the bacterial growth cycle. To make this circuit more stable, we combined the original SLC two-plasmid system into a single operon, one-plasmid system that we then genomically integrated into EcN-lux. In this optimized system, the quorum sensing *plux* promoter drives transcription of both quorum sensing genes, *luxR* and *luxI*, and the phage-derived lysis gene, *ϕX174E* (Fig. 2a). Use of only one promoter and one terminator allowed for this smaller module to be more easily integrated and reduced repeat sequences that can lead to recombination.

**Figure 2:**
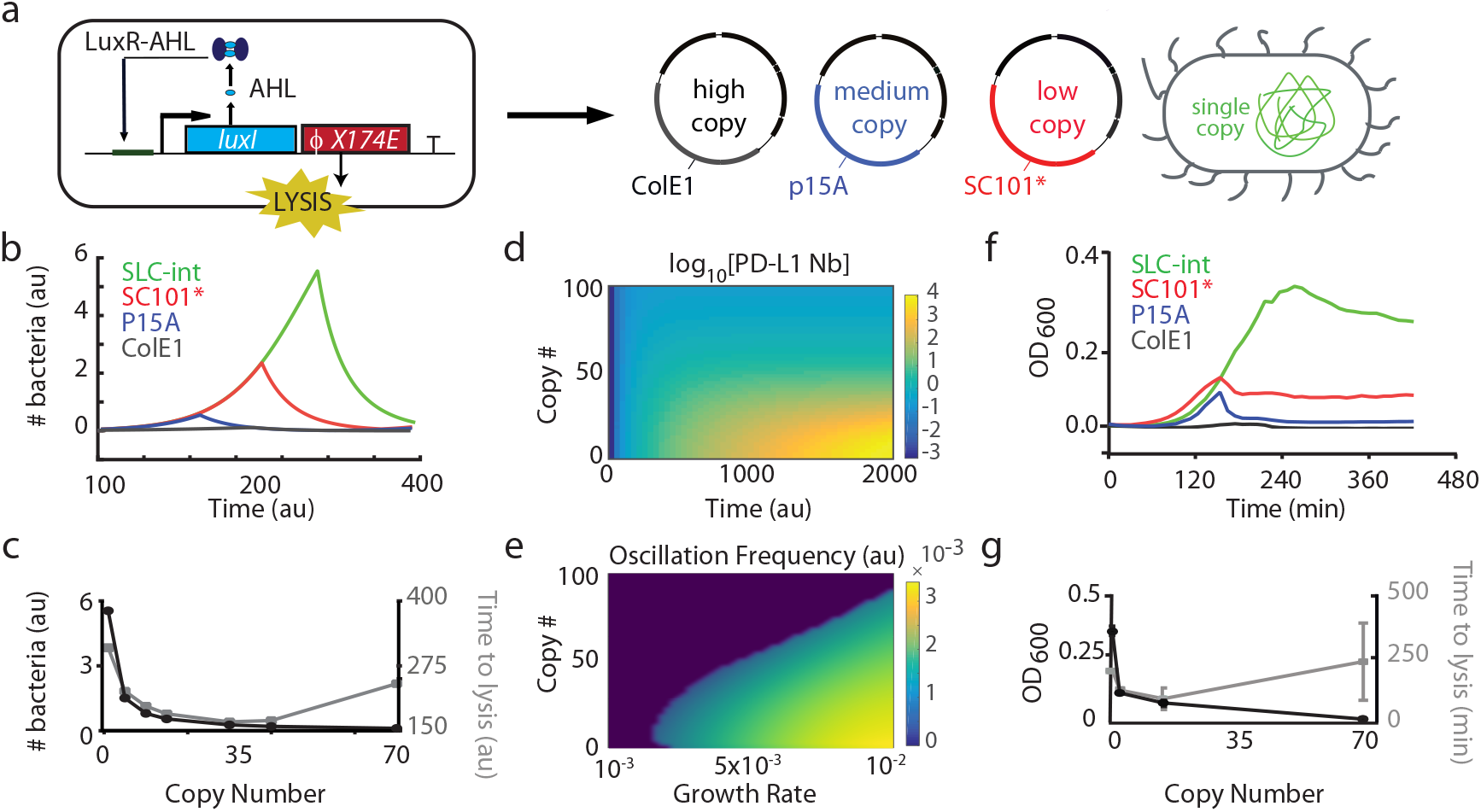
Characterization of lysing variant dynamics. **(a)** Circuit diagram of the single operon SLC circuit in which *plux* drives the transcription of *luxR, luxI* and ϕ 174E genes under a single promoter. The circuit was cloned onto three plasmids of different copy numbers: SC101* (3-4 copies, low), p15A (15-20 copies, medium), and ColE1 (70-100 copies, high) and integrated once into the ϕ80 site of the EcN-lux genome. **(b)** Computational simulation of one lysis event for the copy number variants over time (green: single integrant, red: low copy, blue: medium copy, black: high copy). **(c)** Computational simulation of the number of bacteria needed to reach quorum as a function of copy number (left y axis: black) and time to first lysis event (right y axis: gray). **(d)** Simulated heatmap of the PD-L1 Nb protein produced (z-axis) as a function of both copy number (y-axis) and time (x-axis). **(e)** Simulated heatmap of the oscillation frequency (z-axis) as a function of copy number (y-axis) and growth rate (x-axis). **(f)** Bacterial population dynamics in batch culture of the four lysing variants. **(g)** Experimental data of the number of bacteria needed to reach quorum as a function of copy number and time to first lysis event. Error bars represent S.E.M of repeated experiments

While genomic integration of the SLC operon offers stability, it significantly reduces the copy number of quorum-sensing genes, prompting us to explore how this reduction would affect therapeutic efficacy. Therefore, we computationally modeled the behavior of each circuit variant to further understand the effect of copy number on lysis circuit dynamics. Using a system of ordinary differential equations, we modeled the dynamics of relevant variables in the lysing circuit: *luxI*, ϕ 174E, bacteria number, the small molecule used for bacterial communication AHL (N-Acyl homoserine lactone), and therapeutic production. We first simulated the system to calculate two quantities: number of bacteria required to reach quorum for the first lysis event and time required to reach quorum (Fig. 2b, c). Here, we observed that decreasing copy number of the SLC circuit monotonically increased the number of bacteria required to reach a quorum threshold before lysing. However, time to lysis displayed non-monotonic behavior, where the single copy variant produced a longer time to lysis than medium copy variants, but the high copy variant increased time to lysis again. This non-monotonic behavior is due to the higher level of basal lysis when high copies of the ϕ 174E gene are present, resulting in slower growth, and thus longer time to reach quorum. Further exploration of additional dynamic properties suggested that lower copy variants had a higher mean oscillation amplitude and frequency (Fig. S2a, b).

The system’s behavior at lower copy number variants motivated our exploration into the dynamics of therapeutic production as a direct function of population number over time, where we found that lower copy variants produced more therapeutic in a given period of time and at a faster rate compared to higher copy variants (Fig. 2d, Fig. S2c, d). Furthermore, the average rate of therapeutic production was time invariant, suggesting that this difference in therapeutic production rate was primarily a function of copy number. Given the differences in therapeutic production observed computationally, we sought to understand how our model could be used to predict bacterial dynamics in an *in vivo* setting. Previous studies have shown that bacterial growth kinetics are significantly slower *in vivo*, with the doubling time of the bacteria increasing from ~30 minutes *in vitro* to ~4 hours *in vivo* (26). Using our model, we simulated oscillation frequency as a function of bacterial growth rate and copy number and found that lower copy number variants are more robust to a changing growth rate parameter (Fig. 2e). Together these findings suggest that the SLC-int variant would result in maximal protein production across experimentally-relevant copy numbers.

To experimentally investigate system dynamics of the single operon SLC, we built a library of plasmid variants covering a range of copy numbers for the quorum sensing genes, including a single genomic integration of the operon into the ϕ80 site of the EcN-lux strain (SLC-int) (Fig. 2a). Tracking of bacteria concentration over time in 96 well plates suggested that more copies of the lysing circuit leads to lysis at a lower bacteria concentration with a rapid decay relationship relating these two variables (Fig. 2f, g). Furthermore, the time it took each variant to reach quorum generally suggested that a lower copy number required more time to reach a critical density and the relationship exhibited non-monotonic behavior, consistent with our simulations (Fig. 2g). To additionally visualize lysis events, the SLC-int variant was co-transformed with a constitutive plasmid (ptac:GFP) and imaged over time using fluorescence microscopy (Movie S1). Moreover, multiple oscillations for the SLC-int and SLC-P15A variants were observed by tracking the optical density over time in a plate reader (Fig. S2e).

We next tested whether the SLC-int strain demonstrated lysis behavior and predictable dynamics *in vivo* using a syngeneic CT26 hind flank tumor model (Fig. 3a). We monitored bacteria luminescence from the integrated *luxCDABE* cassette in tumors over time using an In-vivo Imaging System (IVIS) and observed a critical density at approximately 40 hours and population decay to a minimum level at 60 hours (Fig. 3b). Bacteria remaining then repopulated the system until the measurements ceased at the 84 hour time point. When compared to the original SLC two-plasmid system, the SLC-int system took twice as long to reach a critical density, similar to results obtained from our simulations.

**Figure 3:**
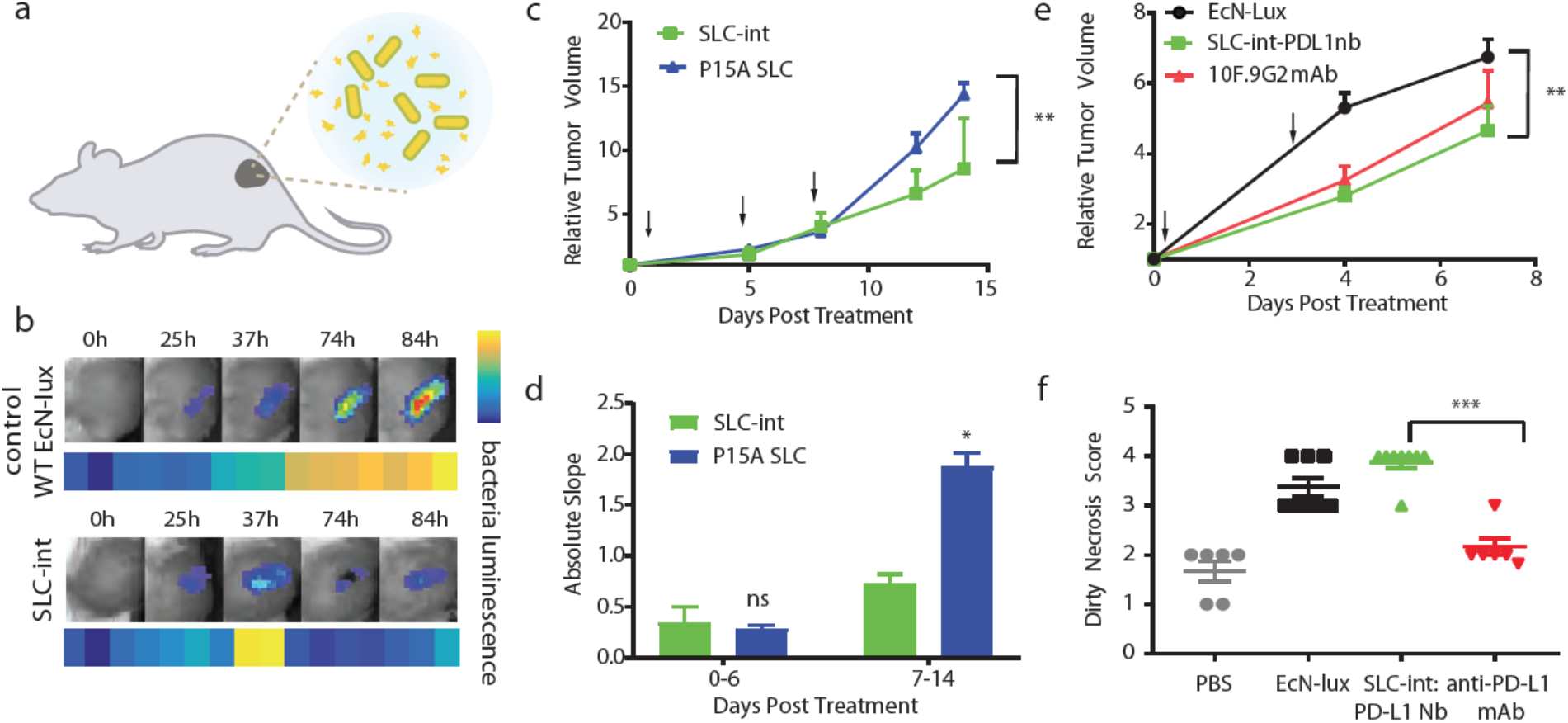
Probiotics expressing PD-L1 Nb (SLC-int:PD-L1nb) elicit a durable therapeutic response in a syngeneic murine model of colorectal cancer. **(a)** BALB/c mice were implanted subcutaneously with 5×10^6^ CT26 cells on both hind flanks and injected with strains genomically expressing a luminescence reporter for more feasible tracking of bacterial population dynamics over time. **(b)** Mice were dosed with 5×10^6^ bacteria of either the non-lysing EcN-lux or SLC-int:PD-L1nb strain and bacterial populations were tracked using IVIS. **(c)** Mean relative tumor trajectory of mice receiving intratumoral injections of 5×10^6^ bacteria of either SLC-int:PD-L1nb or SLC-p15A:PD-L1nb. Arrows indicate days of intratumoral injection (N=6 tumors per group, ** P=0.0065, two-way ANOVA with Bonferroni post-test, error bars represent S.E.M) **(d)** The absolute slope between day 0 and 6 and between day 7 and 14 (* P = 0.0175, segmented linear regression analysis). **(e)** Mean relative tumor trajectory of mice receiving intratumoral injections of 5×10^6^ bacteria of EcN-lux or SLC-int:PD-L1nb or intraperiotoneal injection of 200ug/mouse of 10F.9G2 mAb. All treatments were administered on day 0 and 4 (N=6-8 tumors per group, ** P=0.0015. two-way ANOVA with Bonferroni post-test, error bars represent S.E.M). **(f)** Dirty necrosis scores of histology tissue samples from tumors treated with PBS, EcN-lux, SLC-int:PD-L1nb, and anti-PD-L1 mAb (N=6-8 scores per group of biological replicates, ***P=0.007 Mann Whitney Test ordinal non parametric between SLC-int:PD-L1nb and anti-PD-L1 mAb, error bars represent S.E.M).

### Determining the optimal release mechanism for the PD-L1 Nb

Considering the varying dynamics of the integrated and plasmid-based systems, both the SLC-int and SLC-p15A systems were engineered to produce the PD-L1 Nb. By western blot we were able to confirm production and release of the PD-L1 Nb into the supernatant and observed higher protein levels from the lowest copy SLC-int variant compared to the non-lysing EcN-lux strain transformed with the PD-L1 Nb plasmid at a 16 hour time point (Fig. S3a). We next explored the therapeutic efficacy *in vivo* of both the SLC-int and SLC-p15A copy number variants producing the PD-L1 Nb (SLC-int:PD-L1nb and SLC-p15A:PD-L1nb) in a CT26 hind flank syngeneic mouse model. After 2 weeks, a moderate therapeutic difference was observed, with the SLC-int:PD-L1nb variant resulting in slower relative tumor growth (Fig. 3c). As previously reported, some tumors treated with an anti-PD-L1 monotherapy have a greater response than other tumors within the same treatment group (35, 37). Therefore, we looked at individual mean trajectories and found that for the SLC-int:PD-L1nb treated tumors, 2 of the 6 tumors grew much slower, with 1 tumor trending towards complete regression (Fig. S3b). More interestingly, when therapeutic treatment ceased at day 7 and tumors were left untreated for one week, the relative rate of tumor growth in the SLC-p15A:PD-L1nb variant group increased dramatically in comparison to those in the SLC-int:PD-L1nb group resulting in a significant difference in tumor growth rate (Fig.3d, S3c). This difference in growth rate when treatment stopped is likely to be a result of a more stable SLC-int bacterial system with less plasmid burden. Previous research suggests that increasing bacterial plasmid burden may increase plasmid instability or lead to mutations, which could ultimately affect the therapeutic efficacy of the strain (38). Together these findings led us to choose the SLC-int strain as the release mechanism for the PD-L1 Nb therapeutic.

### Comparison of the SLC-int: PD-L1nb system to anti-PD-L1 mAb

In the syngeneic hind flank mouse model of CT26, comparable therapeutic efficacy was observed for both the 10F.9G2 mAb and the SLC-int:PD-L1nb at day 4 (Fig. 3e). Analysis of individual mean tumor trajectories show that 1 out of 6 tumors in the 10F.9G2 mAb treated group showed minimal relative tumor growth, whereas at least 2 of the 8 tumors in the SLC-int:PD-L1nb treated group showed minimal growth or a reduction in tumor size (Fig. S4). Furthermore, the distribution plot of the absolute tumor volumes for the 10F.9G2 mAb and the SLC-int:PD-L1 nb were similar, with the average absolute tumor volume for the SLC-int:PD-L1 Nb treated group being slightly smaller than that of the 10F.9G2 mAb treated group (Fig. S4). While there was no striking difference in tumor sizes between the two anti-PD-L1 treatment groups, histological analysis indicated a higher dirty necrosis score in bacteria-treated tumors, suggesting that tumors treated with the SLC-int:PD-L1nb had less viable tissue and more neutrophils present in their tumor microenvironment compared to the anti-PD-L1 mAb group (Fig. 3f, Fig. S5).

We explored the versatility of a probiotically-produced PD-L1 Nb by comparing the therapeutic effects of an EcN-lux:SLC-PD-L1nb and the 10F.9G2 mAb in a Balb/c-4T1 syngeneic hind flank mouse model of triple negative breast cancer, where we observed severe toxicity and animal death after only three anti-PD-L1 mAb doses (Fig.S6a, b). After the second dose of the anti-PD-L1 mAb, the body condition of the mice visibly deteriorated with one-third of the mice losing more than 10% of their weight and subsequently remained untreated for the rest of the trial. After the third dose, the mice we had continued treating with the anti-PD-La mAb died and the remaining mice, having never recovered body condition after the second dose, were euthanized (Fig. S6b). In contrast, mice receiving the EcN-lux control and EcN-lux-SLC:PD-L1 nb had no change in body condition and appeared healthy throughout the trial. Toxicity to anti-PD-L1 monotherapy treatment has been previously reported in this breast cancer model (39), but here we have demonstrated that a probiotically-produced PD-L1 Nb mitigates this observed toxicity. Furthermore, our engineered strain demonstrated moderate efficacy when compared to the WT EcN-lux control, suggesting that our engineered probiotic delivery system maintained efficacy while reducing toxicity (Fig. S6c).

### Exploration of immunotherapeutic combination therapies

Due to its local delivery, bacterial therapy may be used to deliver combinations of therapeutics without increasing iRAEs. Therefore, we explored combinations to enhance therapeutic efficacy of our engineered SLC-int:PD-L1nb probiotic therapy in the CT26 model, where only moderate efficacy of the PD-L1 monotherapy was observed consistent with previous results in literature (35, 36). The bacterially-expressed hemolysin exotoxin (hlyE) was chosen for combination with SLC-int:PD-L1nb due to its ability to lyse mammalian cells and potentially release neoantigens that could then be recognized by infiltrating T cells. Furthermore, previous studies have demonstrated moderate efficacy of the hlyE as a monotherapy in mouse models of colorectal cancer (21). The combination of SLC-int:PD-L1nb and EcN-lux producing hlyE (EcN-SLC-hlyE) significantly slowed tumor growth when compared to the individual therapies (Fig. S7a). A distribution plot of individual tumor volumes for each treatment group indicates that tumors with the combination therapy were smaller than those treated with the monotherapies. (Fig. S7b). The overall effect of the combination therapy is more apparent in the individual trajectories, where more mice receiving the combination therapy demonstrated slowed tumor growth over time (Fig. S7c). Lastly, since increased toxicity is observed when immunotherapeutics are used in combination, we tracked body weight of the mice as a proxy for mouse health throughout the trial and saw no significant changes in body weights between treatment groups at the end of the trial (Fig. S7d).

Combination of the SLC:PD-L1nb and EcN-SLC-hlyE therapeutics suggest that the PD-L1 Nb therapeutic platform can be further maximized through combination therapy approaches. Clinically, anti-CTLA-4 mAbs are frequently used in combination with anti-PD-L1 mAbs (40, 41). Therefore, to make our platform more generalizable as a checkpoint blockade treatment option, we engineered a microbial cocktail that could also block the interaction between CTLA-4 and its associated ligands (42, 43). A Nb against CTLA-4 was chosen from the RCSB Protein Data Bank and its sequence was cloned onto the same plasmid vector used to express the PD-L1 Nb (Fig. S8a). In the CT26 mouse model, the SLC-int:CTLA-4nb monotherapy was moderately efficacious and demonstrated similar efficacy to that of the anti-CTLA-4 mAb (Fig. S8b). While the therapeutic response for tumors treated with SLC-int:CTLA-4nb was only modest, less toxicity was observed in these mice compared to the anti-CTLA-4 mAb group, with the mice receiving EcN-lux only and the SLC-int:CTLA-4nb increasing in body weight at a faster rate than those receiving the anti-CTLA-4 mAb. Over the course of two weeks, the average weight of mice treated with the anti-CTLA-4 mAb had increased by ~10%, while the average weight of the SLC-int:CTLA-4nb mice had increased by ~30% (Fig. S8c). Additionally, treatment with the SLC-int:CTLA-4nb induced the production of more IFNγ in the tumor microenvironment when compared to the anti-CTLA-4 mAb and the EcN-lux control (Fig. S8d). An increase in pro-inflammatory cytokines like IFNγ may suggest an increase in T cell infiltration and proliferation, thereby providing further rationale for combining the SLC-int:CTLA-4nb and SLC-int:PD-L1nb therapeutics.

To further explore efficacy in the CT26 model, we treated tumors with a combination of the checkpoint blockade Nbs (Nb cocktail) against PD-L1 and CTLA-4 and observed a significant reduction in the absolute tumor volume compared to the monotherapies (Fig. 4a). Furthermore, mouse body weight was tracked throughout the trial and no significant weight changes were observed (Fig. S9a). To explore the versatility of our combinatorial Nb platform to other models, we tested the Nb cocktail in a syngeneic A20 lymphoma model, where previous literature reports therapeutic efficacy when treated with combination therapies including checkpoint inhibitors (35). We observed a striking efficacy of the Nb cocktail compared to individual SLC-int:CTLA-4nb and SLC-int:PD-L1nb monotherapies, with 3 out of 5 tumors completely regressing and the remaining two tumors showing significant size reduction (Fig. 4b). Moreover, no toxicity was observed and the mice appeared to be healthy and maintain their body condition throughout the trial (Fig. S9b). To further understand the underlying immune response of the observed therapeutic effect, we interrogated the immunophenotype of extracted A20 tumors by flow cytometry and observed a decrease in the the population of regulatory T cells (Tregs, CD4^+^FOXP3^+^) in tumors treated with the Nb cocktail (Fig. 4c). Furthermore, there was an increase in the proliferation of conventional CD4^+^ T cells (CD4^+^FOXP3^−^Ki67^+^) in the Nb cocktail-treated tumors, suggesting that the shift in favor of responsive T cells and decrease in immunosuppressive Tregs may result in a more robust immune response and mediate the therapeutic effect observed (Fig. 4d).

**Figure 4:**
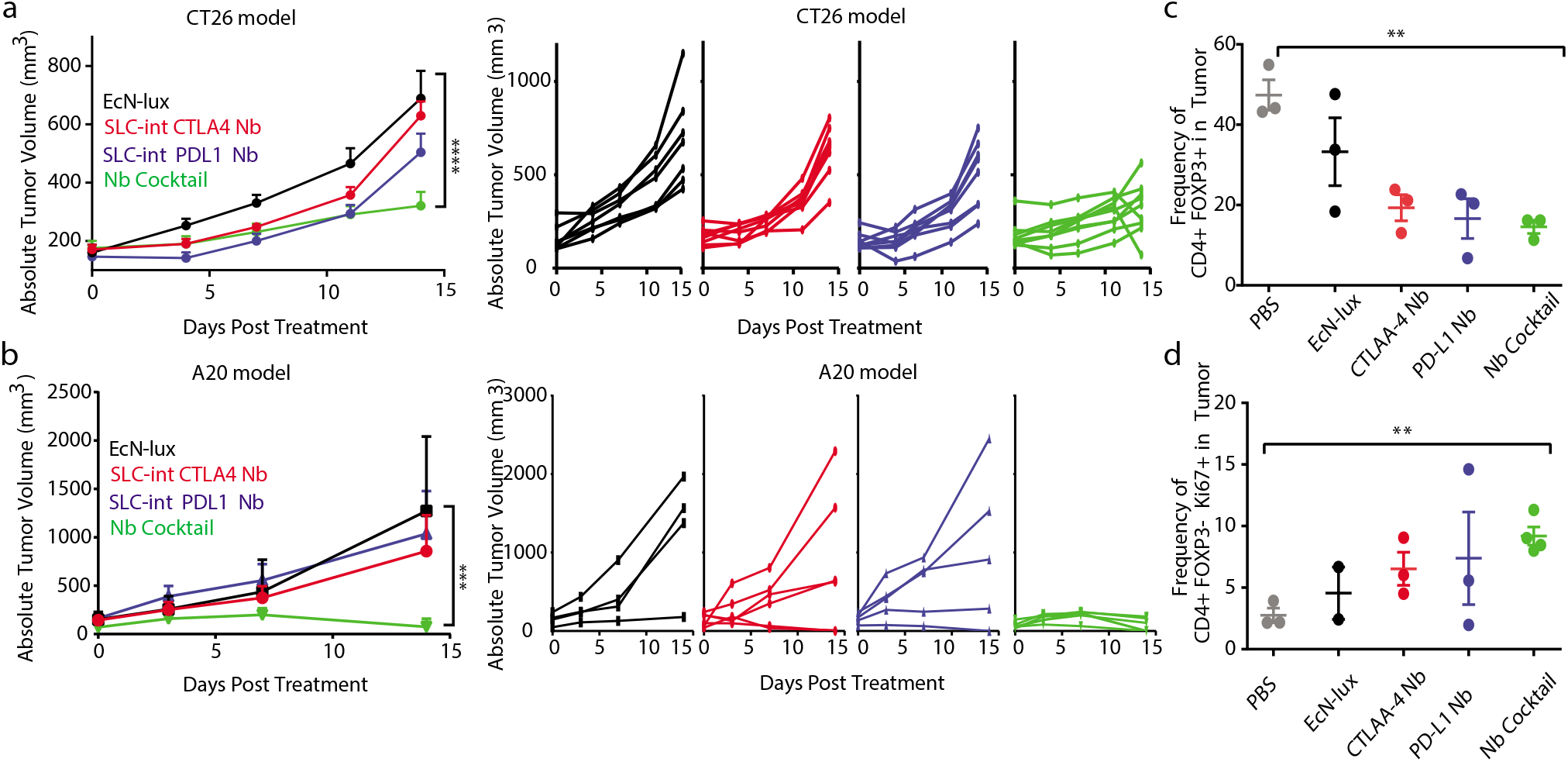
Combination therapy of SLC-int:probiotics producing anti-PD-L1 or anti-CTLA-4 Nbs enhances therapeutic response characterized by a decrease in Treg cells and an increase in proliferative conventional CD4^+^ cells. **(a)** BALB/c mice were implanted subcutaneously with 5×10^6^ CT26 cells on both hind flanks. When tumors reached ~200mm^3^, mice received intratumoral injections every 3-4 days of either EcN-lux, SLC-int:CTLA-4nb, SLC-int:PD-L1nb, or an equal parts Nb cocktail. Mean absolute tumor trajectory and individual trajectories of mice (N=6-8 tumors per group, **** P<0.0001, two way-ANOVA with Bonferroni post-test, error bars represent S.E.M). **(b)** BALB/c mice were implanted subcutaneously with 5×10^6^ A20 cells on both hind flanks. When tumors reached ~150mm^3^ mice received intratumoral injections every 3-4 days of either EcN-lux, SLC-int:CTLA-4nb, SLC-int:PD-L1nb, or an equal parts Nb cocktail. Mean absolute tumor trajectory and individual trajectories of mice (N=4-6, *** P = 0.002, two-way ANOVA, Bonferroni post-test). **(c–d)** Tumor-infiltrating lymphocytes were isolated on day 8 after initial treatment and analyzed by flow cytometry for frequencies of **(c)** CD4^+^FOXP3^+^ regulatory T cells (N=3, unpaired t test between PBS and Nb cocktail, ** P=0.0013) and **(d)** proliferating CD4^+^FOXP3 Ki67^+^ conventional T cells (N=2-3, unpaired t test between PBS and Nb cocktail, ** P = 0.0013). All error bars represent S.E.M.

## DISCUSSION

Here, we have demonstrated the first instance of PD-L1 and CTLA-4 antagonists being expressed by bacteria, which allowed for local therapeutic production and minimized toxicity in multiple syngeneic mouse models. Furthermore, we have characterized novel PD-L1 and CTLA-4 Nbs that can be adapted into other biological circuits and we have optimized their therapeutic release using a genomically-integrated lysing circuit, thereby reducing possible horizontal gene transfer for translational studies. This circuit also serves as a biocontainment measure to confine the bacterial population to the tumor site thereby minimizing the risk of systemic iRAEs.

The SLC-int:Nb systems demonstrated comparable therapeutic efficacy to analogous clinically relevant mAbs and combining the SLC-int:PD-L1 Nb with a bacterially-expressed hemolysin exotoxin led to greater therapeutic efficacy than either monotherapy. Furthermore, the combined Nb cocktail demonstrated tumor shrinkage in the CT26 model and resulted in complete tumor regression in the A20 model. We hypothesize that the synergistic effect of this combination therapy might be a result of fewer immunosuppressive Tregs and an increase in the proliferation of CD4^+^ helper T cells. Moreover, SLC-int:CTLA-4nb led to more detectable IFNγ levels at the tumor site, which could act cooperatively with the SLC-int:PD-L1nb by supporting increased infiltration of immune cells into the tumor microenvironment. Additionally, the production of such cytokines is specifically advantageous for CT26 tumors because colorectal cancers are considered “cold” due to overproduction of prostaglandins that dampen the immune response (44). For this reason, these tumors have only demonstrated modest responses to checkpoint blockade. Therefore, administration of checkpoint blockade nanobodies in vehicles that might lead to more immune cell recruitment could potentially enhance therapeutic responses which would be otherwise difficult to attain for “cold” cancers. To understand the mechanism underlying the therapeutic effects of the SLC-int:Nb system on these types of cancers, further investigation into the immune landscape after treatment will be required.

Cancer immunotherapies are often more effective in combination with other anticancer agents. Therefore, future iterations of the SLC-int system will be programmed to produce a wide variety of immunotherapeutics that can then be tested in combination with the SLC-int:Nb platform. Microbial-based therapeutic platforms are highly modular and optimal for the rapid production of multiple drugs that can then be delivered as a cocktail. Furthermore, to make the system more clinically relevant, other routes of therapeutic administration will be considered. With this in mind, we developed the SLC-int:Nb system in the probiotic strain, *E.coli* Nissle 1917, which has been shown to colonize liver metastases when delivered orally (45), thus offering a more translational route of therapeutic delivery for more advanced metastatic disease.

All together, we have built a stable biological circuit integrated into a probiotic with therapeutics analogous to the current treatment standard for optimization towards clinical translation. The SLC-int:Nb system will help advance the cancer immunotherapy field by providing a delivery vehicle in which combination therapies can be easily explored and toxicities minimized for a broader range of cancer patients.

## MATERIALS AND METHODS

### Study Design

The goal of this study was to develop a probiotic platform that enables the controllable release and effective delivery of checkpoint blockade Nbs within tumors. Due to the preferential growth of microbes within the tumor core, we hypothesized that this delivery platform would reduce toxicities and allow for the delivery of safer and more effective combination therapies. A lysing mechanism was used to release the therapeutic and characterization of these therapeutics, including binding kinetics was established *in vitro* with the CT26 cell line. We then sought to test our delivery system *in vivo*, where all studies were performed on syngeneic hind flank mouse models of CT26, 4T1, or A20 cancers. All mice were randomized prior to treatment and received injections of either monotherapies (SLC-int:PD-L1nb, SLC-hlyE, SLC-int:CTLA-4nb) or combinations of therapies, ensuring that the final concentration of bacteria injected was the same. Caliper measurements were used to track tumor volume, and mouse weight was monitored as a proxy for mouse health. Statistical analysis and sample sizes were determined from previous studies (21, 26, 45). Further details on sample size and replications (technical or biological) are provided in the figure legends.

### Plasmid and Strains

Our circuit strains were cultured in LB media with their respective antibiotics (SC101*variant: 100μg ml^−1^ampicillin, p15A variant: 100μg ml^−1^ spectinomycin, ColE1: 100μg ml^−1^ spectinomycin, and all with 50μg ml^−1^ spectinomycin and kanamycin for strains also transformed with PD-L1 therapeutic plasmid) with 0.2% glucose, in a 37°C shaking incubator. CT26 and 4T1 mammalian cells were purchased from ATCC and cultured in RPMI supplemented with 10% fetal bovine serum and 1% penicillin/streptomycin. A20 cells were purchased from ATCC and cultured in RPMI supplemented with 10% fetal bovine serum and 1% penicillin/streptomycin and 0.01% 2-Mercaptoethanol. Mammalian cells were grown inside a 37°C tissue culture incubator maintained at 5% CO2. Plasmids were constructed using Gibson assembly cloning methods and integrated into the ϕ 80 site of *E.coli* Nissle 1917 using the CRIM protocol (46). The EcN-SLC-hlyE construct was used in previous work from our group (21), while the PD-L1 Nb and CTLA-4 Nb sequence was obtained from an the online RCSB protein data bank and synthesized.

### Antibodies

Antibodies used for *in vivo* experiments include: anti-mouse PD-L1 (BioXCell cat: BE0101and anti-mouse CTLA-4 (BioXCell cat: BE0164). Pre-conjugated primary antibodies used for flow cytometry and microscopy include: Pe/Cy7 anti-mouse CD274 (Biolegend cat: 124313), PE/Cy7 Rat IgG2b (Biolegend cat: 400617), Pe anti-mouse CD274 (Biolegend, cat: 155403), Pe Rat IgG2a isotype (BS biosciences cat: 557076), anti-HA-AF 647 (R&D System cat: IC6875R-025), and mouse IgG_1_-AF 647 (R&D systems cat: IC200R). For western blot an anti-HA high affinity (Roche, cat: 11867423001) and a rat HRP were used. Specific antibodies used for flow cytometry on tumor tissue are listed in the corresponding section.

### Immunofluorescence and Characterization

#### Collecting PD-L1 Nb protein

Non-lysing bacterial strains containing the PD-L1 Nb therapeutic plasmid were grown in a 50mL LB culture with appropriate antibiotics to an optical density of 0.6 then centrifuged at 3000 rcf for 5 minutes. The bacterial pellet was resuspended in 5mL of RPMI media. Samples were frozen at −80°C and thawed in a 30°C incubator 5 times and centrifuged at 3000 rcf for 5 minutes to remove bacterial debris. 1mL of the resulting lysate was then filtered through a 0.2 micron filter.

#### Flow cytometry

For flow cytometry binding experiments, 1×10^6^ CT26 cells were co-incubated in a 14mL round bottom FACS compatible tube with a constant concentration of pre-conjugated PD-L1 Ab (1:800 for the 10F.9G2 clone and 1:200 for the MIH7 clone) and dilutions of the previously prepared bacterial-lysate containing PD-L1 Nb (0.1%, 1%, 10%, 100% of full lysate) in a total working volume of 1mL for 2 hours at room temperature. Samples were spun and washed with ice cold PBS and analyzed on an BD LSRII flow cytometer.

For flow cytometry on *ex vivo* tumor tissue, tumors were extracted for immunophenotyping on day 8 following bacteria treatment on days 0, 4, and 7. Lymphocytes were isolated from tumor tissue by mechanical homogenization and digestion with collagenase A (1mg/ml, Roche) and DNAase I (0.5ug/mL, Roche) in isolation buffer (RPMI 1640 supplemented with 5% FBS, 1%L-glutamine, 1%penicillin/streptomycin, and 10mM Hepes) for 1h at 37C. Cells were then filtered through 100um cell strainers and washed in isolation buffer before staining. A Ghost Dye cell viability stainin was used as a live/dead marker. Extracellular antibodies used include: anti-B220 (BD 1:400), anti-CD4 (Tonbo), anti-CD8 (eBioscience 1:400) and anti-NKp46(BD 1:200). Cells were then fixed using FOXP3/transcription factor staining buffer set (Tonbo) as per the manufacturer’s protocol and then stained intracellularly. Intracellular antibodies used include: anti-TCRβ(BD 1:400), anti-Ki67(Thermo 1:400) and anti-FOXP3(ebioscience 1:400). Samples were analyzed using a BD LSR Fortessa cell analyzer.

Single-stains and isotype controls were used for compensation of the dual stain. FlowJo was used for all data analysis.

#### Microscopy

For microscopy, 1×10^6^ CT26 cells were stained on a poly-l-lysine coated glass flat bottom plate with appropriate antibodies for detection of PD-L1 expression (10F.9G2 clone). In a separate experiment, full bacterial lysate containing PD-L1 Nb was incubated on the monolayer of CT26 cells and probed with a pre-conjugated anti-HA antibody. A Nikon Eclipse Ti microscope was used for all microscopy and FIJI was used for any post-image analysis.

### Characterization of PD-L1 Nb Expression and Release

To determine protein expression and functionality of the HA tag, PD-L1 Nb protein was released using the freeze/thaw method described above. A western blot was use for protein visualization and was probed for 1hr with the primary anti-HA antibody (1:800) and 1 hr with a rat HRP (1:4000) at room temperature. Protein was detected using a chemiluminescent substrate.

### *In vivo* Studies

#### Tumor models

Animal experiments were performed on 6-8 week old female BALB/C mice from Taconic with bilateral subcutaneous hind flank tumors from an implanted mouse colorectal cancer cell line CT26, mouse lymphoma cell line A20, or a mouse breast cancer cell line, 4T1. Tumor cells were prepared for implantation at a concentration of 5e7 cells/ml in RPMI without phenol red. Cells were implanted at 100uL per flank, with each implant consisting of 5×n10^6^ cells. Tumors grew for approximately 10 days or until the diameter was 4-10mm in diameter or volume was ~200mm^3^ for CT26 and 4T1 tumors and ~150mm^3^ for A20 tumors. All mice were randomized prior to treatment.

#### Bacteria growth and therapeutic administration

Bacterial strains were grown overnight in LB media containing appropriate antibiotics and 0.2% glucose for less than 12 hours. The overnight was subcultured at a 1:100 dilution in 50mL of fresh media with antibiotics and glucose and grown until an OD of approximately 0.05 to prevent bacteria containing SLC operon from reaching quorum. Bacteria were spun down at 3000 rcf and washed 3 times with sterile ice-cold PBS. Bacteria were delivered intratumorally at a concentration of 5e7 cells/ml in PBS with a total of 30-40uL injected per flank every 3-4 days (OD of 0.1 is equivalent to 1e8 cells/mL). The PD-L1 mAb and the CTLA-4 mAb were injected intraperitoneally at a concentration of 200ug/mouse and 100ug/mouse respectively.

#### Tumor and bacteria monitoring in vivo

CT26 and 4T1 tumor volumes were quantified using calipers to measure length, width, and height of each tumor with volumes being reported as LxWxH. A20 tumors were measured as (LxW)/2. Tumors were measured every 3 to 4 days and mouse weight was tracked to monitor overall mouse health. All bacterial strains used were luminescent, which could be measured with the In Vivo Imaging System. Living Image software was used for luminescent quantification.

#### Histology

Tumors were extracted, fixed in 10% formalin, and sent to the Histology and Imaging Core at the University of Washington, where the tissue was processed and Haemotoxylin and Eosin stained. The pathologist blinded to sample treatments manually scored the samples for dirty necrosis (0 = not present, 1 = minimal, 2 = mild, 3 = moderate, and 4 = severe, abscess like)

#### Cytokine Analysis

Tumors were extracted from mice on day 14 following initial treatment. After one snap-freeze cycle, tumors were mechanically lysed in complete lysis buffer (1x Protease Inhibitor cocktail, Sigma Aldrich Cat: P27using a gentleMACS Dissociator.14-1BTL; EMD Millipore lysis buffer (Cat: 43-040) using a gentleMACS tissue dissociate. Once fully dissociated, the tumor homogenates were centrifuged for 10min at 10,000g to clear debris. The lysate was spun for 5min at 5,000g. Pre-cleared lysates were stored at −80C until analysis by Luminex Multiplex Assay. A Bradford colorimetric assay was used to normalize protein concentrations and 35ug lysate in 25uL volume was analyzed in the Luminex Multiplex Assay (EMD Millipore) according to the manufacturer’s protocol.

#### Statistical Analysis

Statistical tests were calculated in GraphPad Prism 7.0. The details of the statistical tests are indicated in the respective figure legends. Where data was assumed to be normally distributed, values were compared using a one-way ANOVA for single variable or a two-way ANOVA for more than one variable with a Bonferroni post-test applied for multiple comparisons. For categorical data comparisons, data was assumed to be nonparametric and a Mann Whitney U Rank test was used for single variable, two group comparisons.

## SUPPLEMENTARY MATERIALS

Figure S1: Engineering and characterization of the ptac: PD-L1 Nb plasmid.

Figure S2: Computational modeling of the lysing variant dynamics.

Figure S3: Characterization of lysing circuit variants in a syngeneic colorectal model.

Figure S4: Individual tumor trajectories for comparison between probiotic expressing PD-L1nb (SLC-int:PD-L1nb) and systemically delivered antibody in CT26 model.

Figure S5: Histological images and dirty necrosis scoring in CT26 mouse tumors.

Figure S6: Comparison between probiotically-produced PD-L1 Nb and systemically delivered antibody in 4T1 model.

Figure S7: SLC-int:PD-l1nb and bacterially-expressed hemolysin exotoxin (hlyE) combination therapy is more efficacious than individual therapies.

Figure S8: Engineered probiotic expressing CTLA-4 Nb.

Figure S9: Relative body weight for combination of PD-L1 and CTLA-4 probiotics.

Mathematical Model

Movie S1: Visualization of SLC-int lysis.

## ACKNOWLEDGEMENTS

This work was supported in part by the NIH Pathway to Independence Award (R00CA197649-02), DoD Idea Development Award (LC160314), DoD Era of Hope Scholar Award (BC160541), and the National Science Foundation Graduate Research Fellowship under Grant No. (1644869). Research reported in this publication performed in the CCTI Flow Cytometry Core, was supported in part by the Office of the Director, National Institutes of Health under awards S10RR027050. Additionally, these studies used the resources of the Cancer Center Flow Core Facility funded in part through Center Grant P30CA013696. The content is solely the responsibility of the authors and does not necessarily represent the official views of the National Institutes of Health. We would like to thank Omar Din for assistance with the computational modeling and *in vitro* lysing circuit variant dynamics. We would also like to thank members of the Danino lab for critical review of the manuscript.

## AUTHOR CONTRIBUTIONS

C.G. and T.D. conceived and designed the therapeutic platform. C.G. built and characterized the library of SLC variants, optimized the computational modeling platform, and performed experiments *in vitro* to test bacterial strains and characterize the immunotherapeutic. C.G., C.C., T.H., I.L, and S.C. designed and performed *in vivo* experiments. C.G and N.A designed and analyzed the immune characterization assays. P.T performed histology. C.G. and T.D. analyzed data and wrote the manuscript with input from all of the other authors.

## DATA AVAILABILITY

The data that support the findings of this study are available within the paper and its supplementary information files. Additional data are available from the authors upon reasonable request.

## SUPPLEMENTARY MATERIALS

**Figure S1:**
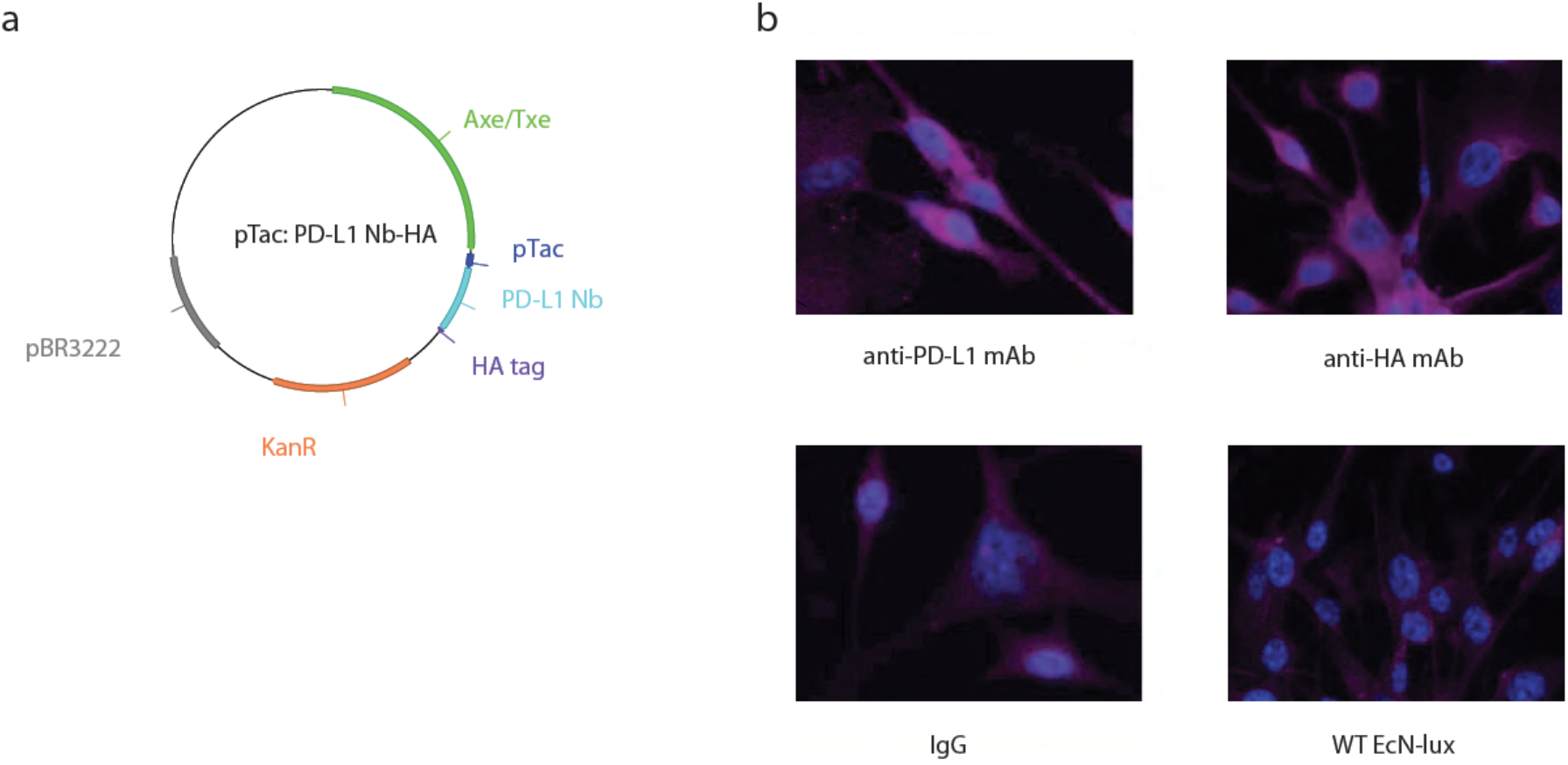
Engineering and characterization of the ptac: PD-L1 Nb plasmid. **(a)** Plasmid map for the constitutive expression of the PD-L1 Nb on a high copy plasmid. **(b)** Immunofluorescence microscopy image of CT26 cells (purple: PD-L1, blue: nuclei). Left panel shows CT26 cells stained for PD-L1 expression with a flourescently-tagged 10F.9G2 mAb and its respective isotype. Right panel shows CT26 incubated with bacterial lysate containing PD-L1 Nb where the PD-L1 Nb is probed for using an anti-HA mAb.

**Figure S2:**
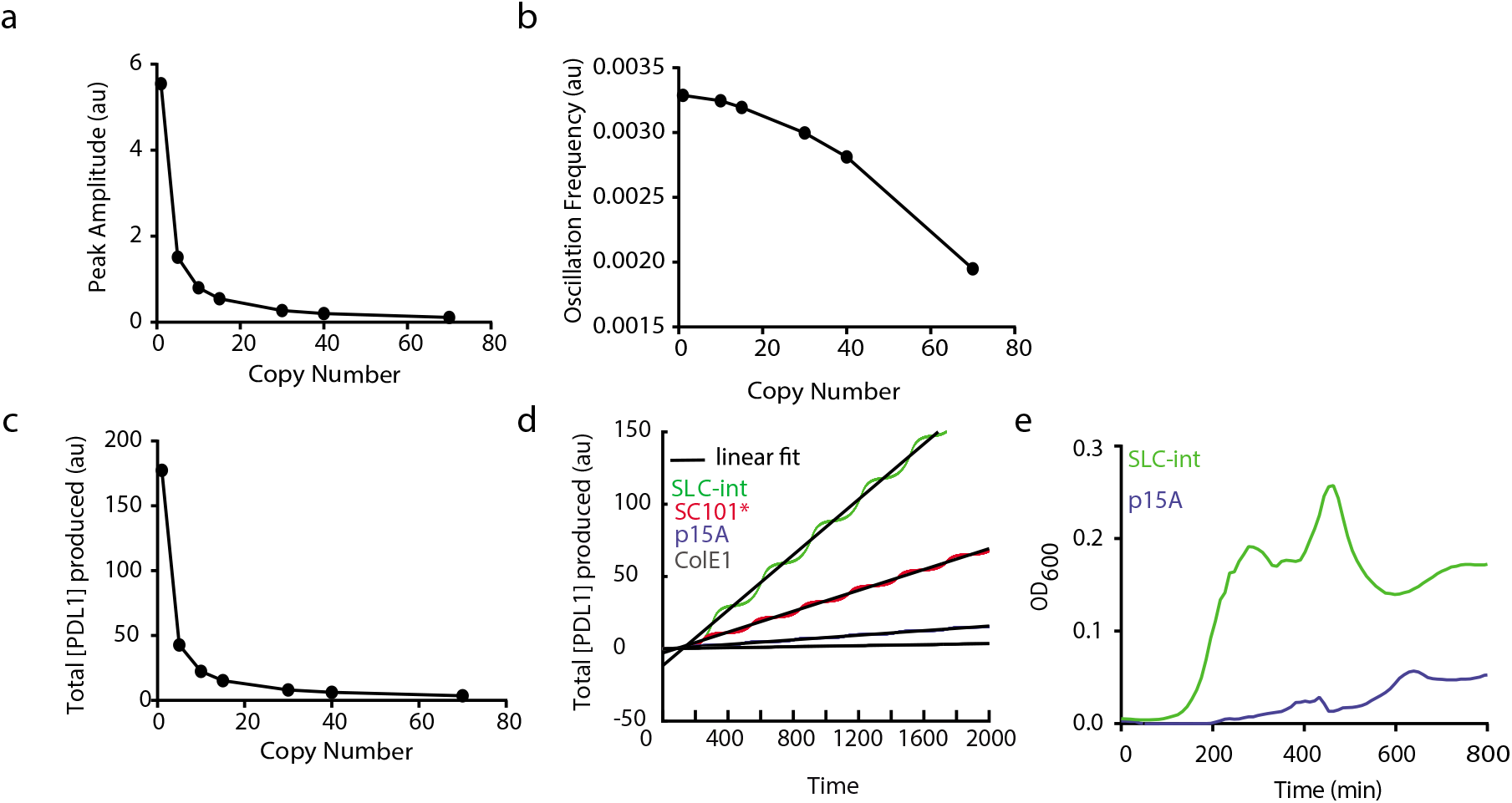
Computational modeling of the lysing variant dynamics. **(a)** Peak amplitude of bacterial population at quorum as a function of copy number. **(b)** Frequency of oscillations over a given time frame as a function of copy number. **(c)** Total PD-L1 Nb produced over a given time frame as a function of copy number. **(d)** Rate of PD-L1 Nb production for copy number variants. **(e)** Experimental dynamics measured in a Tecan plate reader for SLC-int and p15A copy number variants showing multiple lysis events.

**Figure S3:**
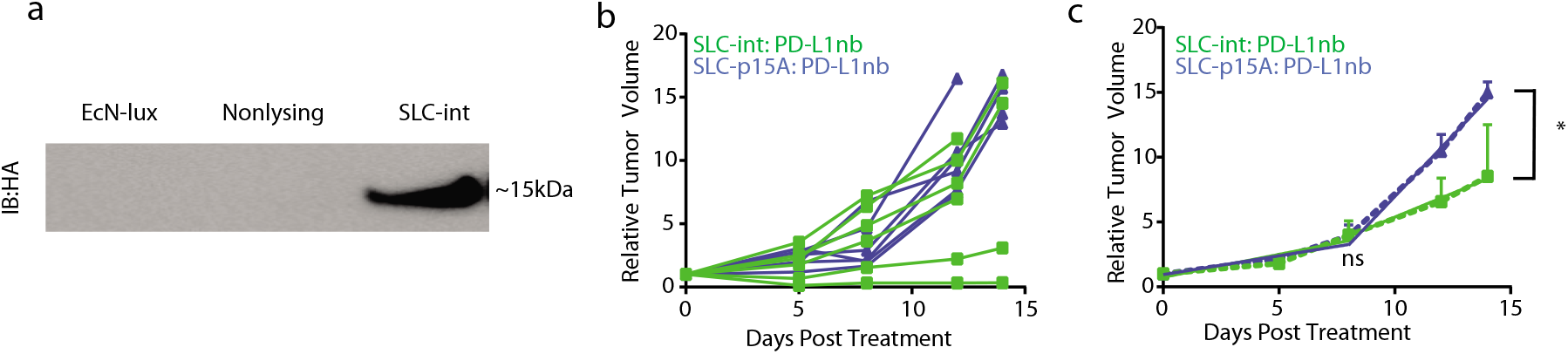
Characterization of lysing circuit variants in a syngeneic colorectal model. **(a)** Western blot showing PD-L1 Nb protein levels collected from the supernatant of EcN-lux, EcN-lux:PD-L1nb (non-lysing), and SLC-int:PD-L1nb strains. **(b)** Individual trajectories of mean tumor volume for CT26 tumor bearing BALB/c mice receiving injections of the SLC-int:PD-L1nb or SLC-p15A: PD-L1nb variant on day 0, 4 and 7 post treatment. **(c)** Segmented linear regression (dotted line) analysis of the relative tumor volume from 0-6 and 7-14 days. For days 0-6, linear regression fit R^2^ = 0.85 and 0.99 for SLC-int:PD-L1nb and SLC-p15A:PD-L1nb groups respectively. For days 7-14, linear regression fit R^2^ = 0.99 and 0.99 for SLC-int:PD-L1nb and SLC-p15A:PD-L1nb groups respectively. N=6 per group, *P = 0.0175, error bars represent S.E.M.

**Figure S4:**
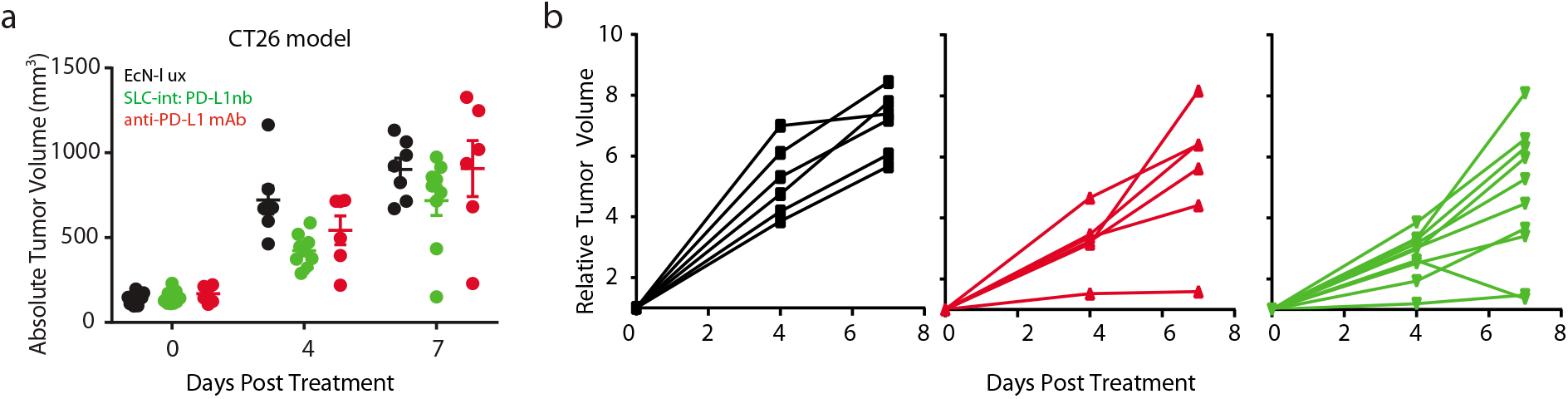
Individual tumor trajectories for comparison between probiotic expressing PD-L1nb (SLC-int:PD-L1nb) and systemically delivered antibody in CT26 model. **(a)** Plot showing distribution of tumor sizes in CT26 model. **(b)** Individual relative tumor volumes for CT26 tumor bearing BALB/c mice injected with EcN-lux, SLC-int:PD-L1 Nb, and 10F.9G2 mAb. This experiment corresponds to Figure 3e in the text.

**Figure S5:**
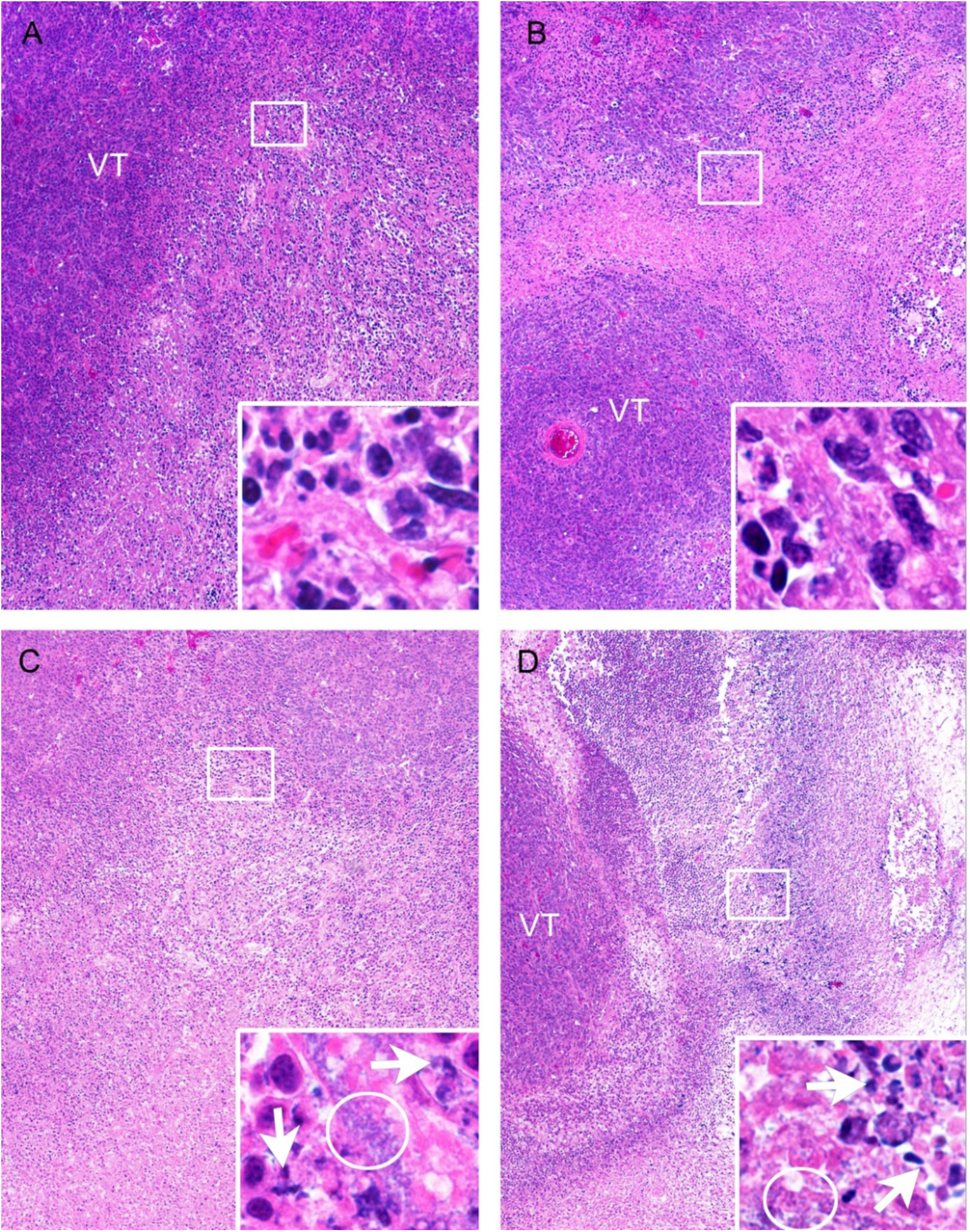
Histological images and dirty necrosis scoring in CT26 mouse tumors. Main panels are 40x, inserts are 600x of the approximate areas of boxed regions in low magnification images. VT is viable tumor; Circles in insets are bacterial colonies; arrows degenerate neutrophils (i.e. dirty necrosis). **(A)** PBS, **(B)** 10F.9G2 mAb, **(C)** EcN-lux, **(D)** SLC-int:PD-L1-nb. This experiment corresponds to Figure 3e in the text.

**Figure S6:**
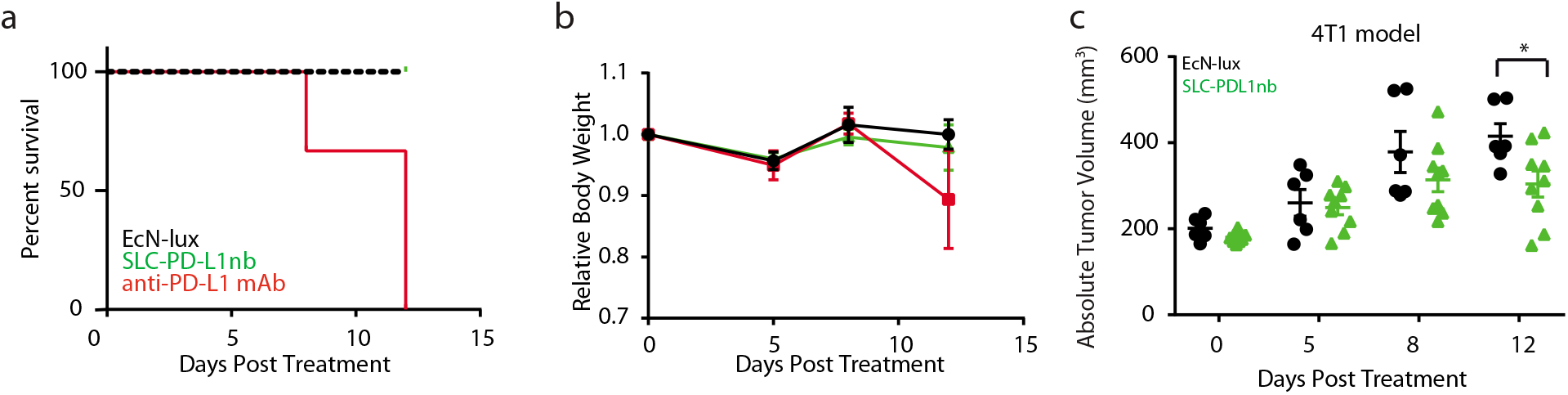
Comparison between probiotically-produced PD-L1 Nb and systemically delivered antibody in 4T1 model. BALB/c mice were implanted subcutaneously with 5×10^6^ 4T1 cells on both hind flanks. When tumors reached ~200mm^3^, mice received intratumoral injections of 5×10^6^ bacteria every 3-4 days of either EcN-lux, SLC-PD-L1nb, or 200ug/mouse of 10F.9G2 mAb **(a)** Survival of mice following 5×10^6^ bacteria dosed on days 0, 5, and 8 days post treatment, and systemically delivery mAb dosed on days 0, 5, 8 and 12. Severe toxicities in weight and body conditions of antibody treated mice were observed, leading to death or required euthanasia of all mice within two weeks. **(b)** Mean relative body weight. **(c)** Distribution of tumor sizes. N=6-8 per group, *P = 0.0199, two-way ANOVA with Bonferroni post-test, error bars represent S.E.M.

**Figure S7:**
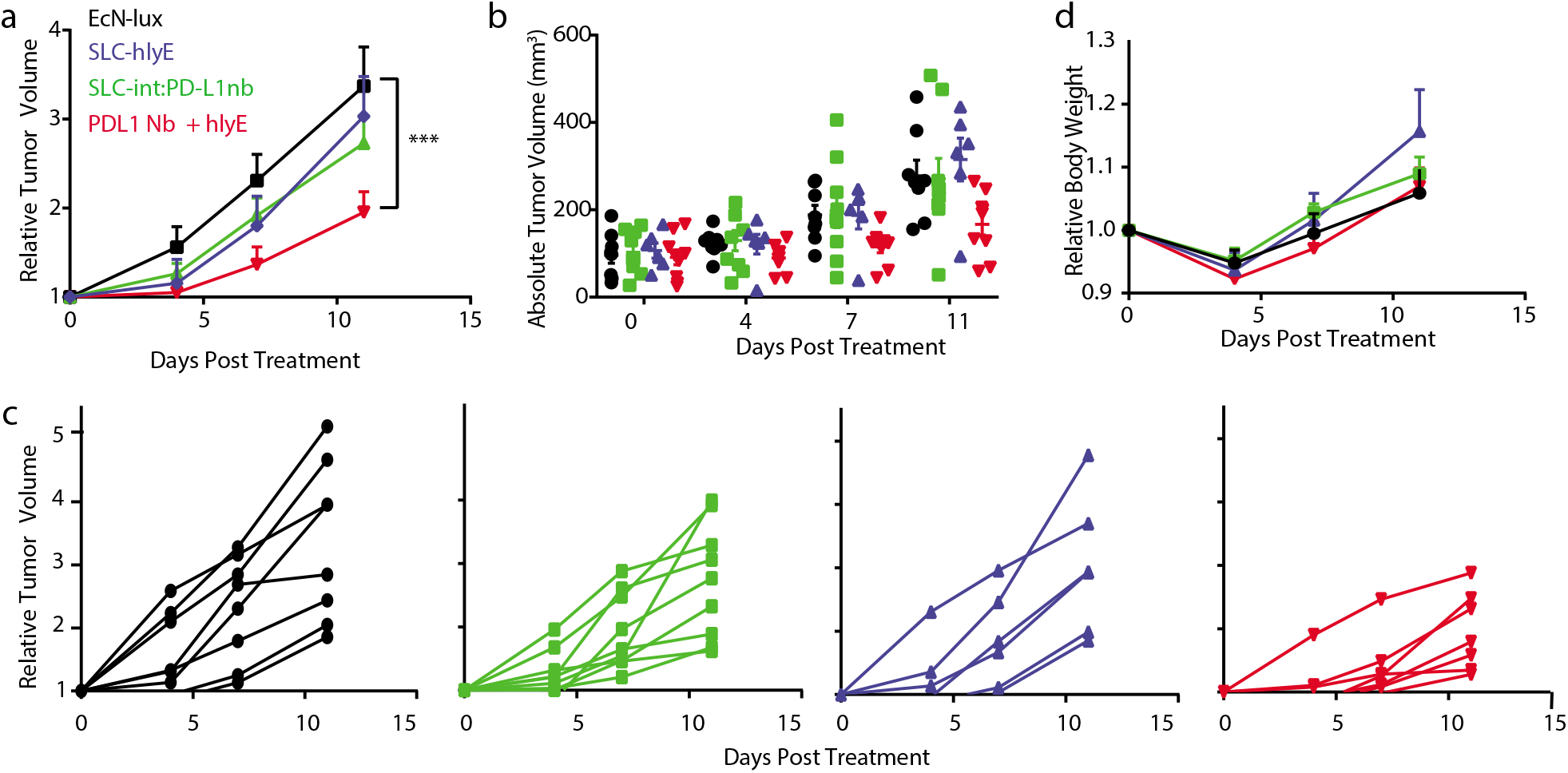
SLC-int:PD-l1nb and bacterially-expressed hemolysin exotoxin (hlyE) combination therapy is more efficacious than individual therapies. BALB/c mice were implanted subcutaneously with 5×10^6^ CT26 cells on both hind flanks. When tumors reached ~200mm^3^, mice received intratumoral injections of 5×10^6^ bacteria every 3-4 days of either EcN-lux, EcN-SLC-hlyE, SLC-int:PD-L1nb, or an equal parts cocktail of EcN-SLC-hlyE and SLC-int:PD-L1nb strains. (a) Mean relative tumor trajectory of mice. **(b)** Distribution of tumor sizes. **(c)** Individual trajectories of relative tumor volumes for each treatment group. N=6-9 tumors per group, *** P=0.0004, twoway ANOVA with Bonferroni post-test, error bars represent S.E.M. **(d)** Mean relative body weight.

**Figure S8:**
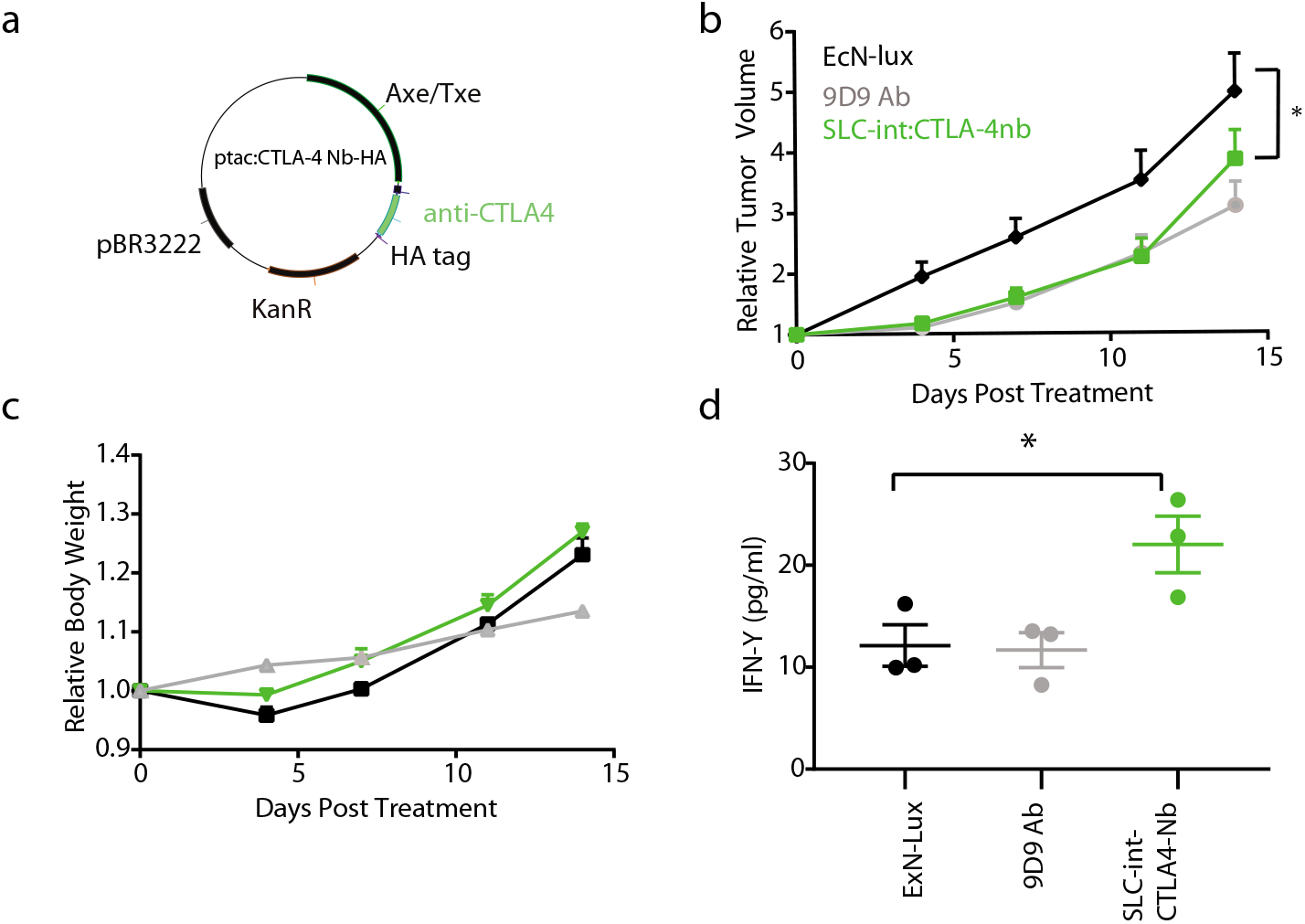
Engineered probiotic expressing CTLA-4 Nb. **(a)** Plasmid map for the constitutive expression of the CTLA-4 Nb on a high copy plasmid. **(b)** BALB/c mice were implanted subcutaneously with 5×10^6^ CT26 cells on both hind flanks. When tumors reached ~200mm^3^, mice received intratumoral injections every 3-4 days of 5×10^6^ bacteria of either EcN-lux, SLC-int:CTLA-4nb or 100ug/mouse intraperitoneal injections of a 9D9 mAb. Graph shows mean relative tumor trajectories. **(c)** Mean relative body weight. N=6-7 per group, *P=0.0407, two-way ANOVA with Bonferroni post-test, error bars represent S.E.M. **(d)** Cytokine levels of tumor lysates measured by Luminex Multiplex Assay. N=3 biological replicates per group, *P=0.0224, one-way ANOVA with Bonferroni post-test, error bars represent S.E.M.

**Figure S9:**
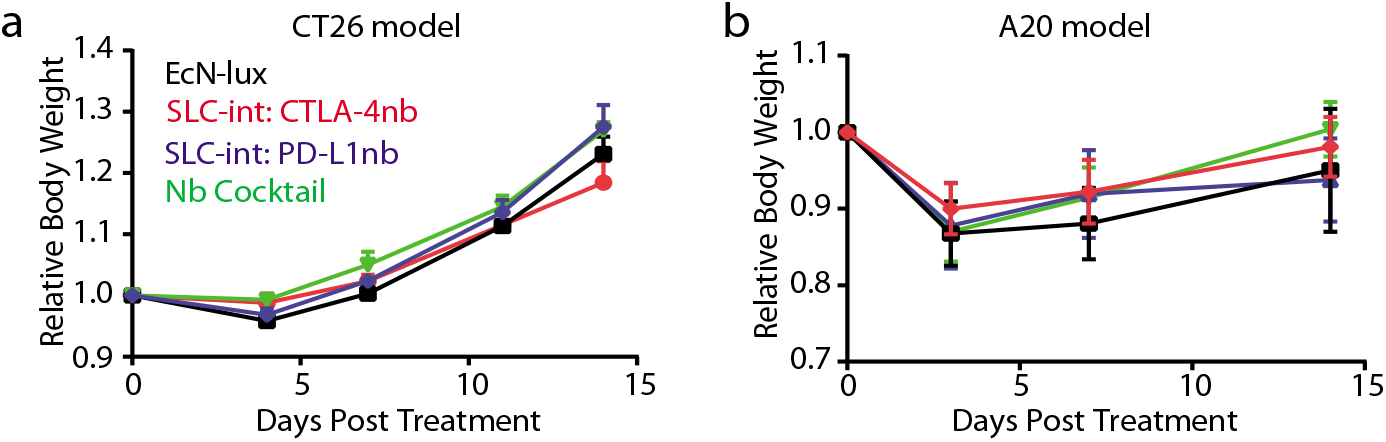
Relative body weight for combination of PD-L1 and CTLA-4 probiotics. Mean relative body weight in a syngeneic **(a)** CT26 and **(b)** A20 mouse model. This experiment corresponds to Figure 4 in the text.

## Mathematical Model

We built upon the ordinary differential equation model explored in the original sycnhrionized lysis circuit system (19). The following set of ordinary differential equations were derived to describe the bacterial population number (N, equation[1]), total extracellular AHL (H, equation[2]), intracellular concentrations of the lysis protein E (L, equation[3]), intracellular concentrations of the LuxI protein (I, equation[4]), and intracellular concentrations of the PD-L1 Nb protein (P, equation[5]). In this system, bacteria grow logistically at a rate of μ_N_ to a maximum capacity, N_0_ and lyse once at quorum. AHL is produced at a rate proportional to bacterial population and is cleared out at a rate of μ. The lysis protein is produced at a rate proportional to copy number and has a basal degradation (γ_L_) and is also diluted as the bacterial population grows (μ_g_). The luxI protein has similar dynamics, but is further degraded by ClpXP machinery (γ_c_). The therapeutic protein production is proportional to bacteria number and has basal degradation (γ_p_) and is diluted as the bacterial population grows (μ_g_). Internal production of LuxI and lysis protein is described by P_lux_ (equation[6]) and the rate of cell degradation due to lysis is described by the hill function, γ_N_ (equation[7]).

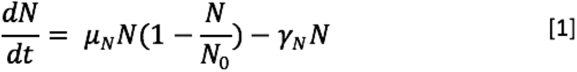

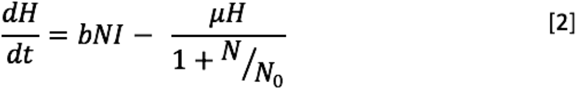

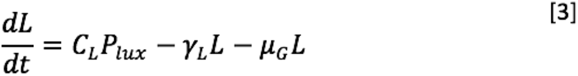

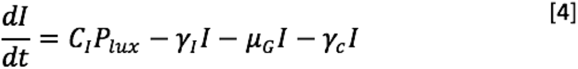

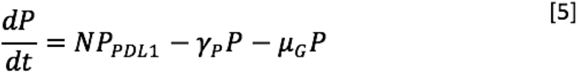

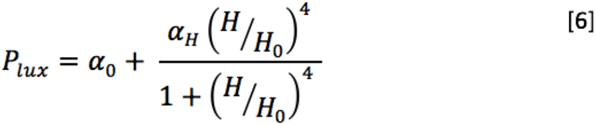

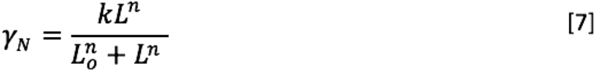

### Model Parameters Values

To explore the dynamics of our library of lysing variants, we iterated through a large range of copy numbers, *C_I_* and *C_L_* ranging from 1 to 100. Growth rate (μ_N_). was experimentally-derived and the doubling time of ~30 minutes for SLC-int:PD-L1nb was used to calculate a growth rate of 0.023min^−1^ for *in vitro* plots (Fig. 2b, c, Fig. S2a-d) using the doubling time formula. Based on previous literature (24), the doubling time of bacteria increases to ~4 hours, which yielded a growth rate of 0.00289min^−1^ (Fig. 2d, e). Since growth rate and dilution due to cell growth was assumed to be equal μg was set equal to μN. Other parameter values include: α_0_ = 0.1, α_H_ = 50, h_0_=5, b=20, μ_g_ = 1.5, L_0_ = 8, n=2, m=4. γ_L_ = 1.5, K =0.05, γ_I_ = 5, γ_C_ = 12, N_0_=10.

**Movie S1: Visualization of SLC-int lysis.** The EcN-lux:SLC-int strain was transformed with a ptac:sfGFP plasmid. Bacteria were seeded onto a 96 well glass bottom plate and timelapse was used to capture GFP fluorescence change over time at 10X magnification.

